# Establishment and application of unbiased *in vitro* drug screening assays for the identification of compounds against *Echinococcus granulosus s.s*

**DOI:** 10.1101/2023.05.02.539024

**Authors:** Marc Kaethner, Matías Preza, Tobias Kaempfer, Pascal Zumstein, Claudia Tamponi, Antonio Varcasia, Andrew Hemphill, Klaus Brehm, Britta Lundström-Stadelmann

## Abstract

*Echinococcus multilocularis* and *E. granulosus s.l.* are the causative agents of alveolar and cystic echinococcosis, respectively. Drug treatment options for these severe and neglected diseases are limited to benzimidazoles, which are not always efficacious, and adverse side effects are reported. Thus, novel and improved treatments are needed.

In this study, the previously established platform for *E. multilocularis in vitro* drug assessment was adapted to *E. granulosus s.s.*. In a first step, *in vitro* culture protocols for *E. granulosus s.s.* were established. This resulted in the generation of large amounts of *E. granulosus s.s.* metacestode vesicles as well as germinal layer (GL) cells. *In vitro* culture of these cells formed metacestode vesicles displaying structural characteristics of metacestode vesicles generated *in vivo*. Next, drug susceptibilities of *E. multilocularis* and *E. granulosus s.s.* protoscoleces, metacestode vesicles and GL cells were comparatively assessed employing established assays including (i) metacestode vesicle damage marker release assay, (ii) metacestode vesicle viability assay, (iii) GL cell viability assay, and (iv) protoscolex motility assay. The standard drugs albendazole, buparvaquone, mefloquine, MMV665807, monepantel, niclosamide and nitazoxanide were included. MMV665807, niclosamide and nitazoxanide were active against the parasite in all four assays against both species. MMV665807 and monepantel were significantly more active against *E. multilocularis* metacestode vesicles, while albendazole and nitazoxanide were significantly more active against *E. multilocularis* GL cells. Albendazole displayed activity against *E. multilocularis* GL cells, but no effects were seen in albendazole-treated *E. granulosus s.s.* GL cells within five days. Treatment of protoscoleces with albendazole and monepantel had no impact on motility. Similar results were observed for both species with praziquantel and its enantiomers against protoscoleces. In conclusion, *in vitro* culture techniques and drug screening methods previously established for *E. multilocularis* were successfully implemented for *E. granulosus s.s.,* allowing comparisons of drug efficacy between the two species.

## Introduction

Alveolar echinococcosis (AE) and cystic echinococcosis (CE) are severe zoonotic diseases caused by the tapeworms *Echinococcus multilocularis* and *E. granulosus sensu lato* (*s.l*.), respectively. Both parasitoses are acquired via oral uptake of eggs containing infectious oncospheres [1]. Subsequently, these oncospheres invade the liver and other organs, and establish the disease-causing stage, the *Echinococcus* metacestode. Natural intermediate hosts of *E. multilocularis* are rodents and other small mammals, whereas *E. granulosus s.l.* naturally infects a variety of larger mammals such as sheep, cattle, and others, depending on parasite genotype. Inside metacestodes, protoscoleces develop that will differentiate into adult worms when ingested by definitive hosts such as foxes or dogs [1]. *E. multilocularis* and *E. granulosus s.l.* can also infect aberrant hosts that usually do not further transmit the parasite, but still acquire the diseases AE or CE. Both parasites thereby impose considerable burden on human and veterinary health and are responsible for high economic losses, in particular in the case of CE [2]. On a global scale, *E. multilocularis* and *E. granulosus s.l.* are the third and second most important food-borne parasites, respectively [3]. AE and CE cause global burdens of 688,000 and 184,000 Disability Adjusted Life Years (DALY), and are responsible for at least 18,500 and 188,000 new human cases per year [4]. In Europe, they are the first and fourth most important food-borne parasites [5,3]. However, the actual number of echinococcosis cases may be significantly higher as not all infections are officially recorded [6–8]. In addition, there is a reported emergence of AE in Europe and Canada [9,10], of CE in the Middle East [11], and both AE and CE in the East Asia-Pacific region and Central Asia [12,13].

The disease-causing stage of *E. multilocularis* and *E. granulosus s.l.* is the metacestode, which grows as a multivesicular (AE) or unilocular (CE) cyst filled with vesicle fluid. Each metacestode vesicle is surrounded by an outer acellular and carbohydrate-rich laminated layer. This layer is followed by an inner syncytial tegument and a germinal layer (GL) [14,15], all of which build up the metacestode tissue. The GL contains a variety of differentiated cells including muscle cells, nerve cells, subtegumentary cytons, glycogen storage cells and undifferentiated stem cells [16,17]. Especially the latter are responsible for the high regenerative potential of this parasite.

In the human host, *E. multilocularis* metacestodes mainly infect the liver and subsequent proliferation leads to infiltration of the affected organ [1]. If feasible, patients should undergo radical surgery, which can result in curative treatment if all parasitic tissue is removed, but at least two years of post-surgical chemotherapy is recommended. The alternative option is lifelong chemotherapy with daily treatment of 10 to 15 mg/kg albendazole (ABZ) or 40 to 50 mg/kg mebendazole, both exhibiting limited efficacy [18].Treatment is not effective in all cases, and the daily treatment with benzimidazoles can cause adverse side effects, most notably hepatotoxicity, and this normally leads to treatment discontinuation and, as a consequence, recurrence of the disease [19,20]. This recurrence is most likely caused by stem cells in the GL [21,16], which might express an isoform of the benzimidazole target beta-tubulin that is not efficiently bound by ABZ or mebendazole [22]. Thus, it has been suggested that GL cells survive benzimidazole treatment, which then leads to parasite regrowth [22]. Since the currently applied drug treatment does not act parasiticidally and AE causes mortality in case the treatment fails with no alternative treatment options in sight, there is a strong need for the development of new drugs that can target also the stem cells of the parasite [23,24].

*E. multilocularis* represents a model cestode, due to the fact that *in vitro* culture of metacestode vesicles and isolated GL cells (including parasite stem cells) has been established, and options for genetic manipulation by RNAi were introduced [25,26]. In addition, parasite isolates can be passaged in mice and cryopreservation protocols are implemented [27–30]. While older studies on the evaluation of drug effects against *E. multilocularis* relied on subjective microscopic assessments, the more advanced culture techniques for *E. multilocularis* enabled researchers to develop a drug screening cascade for the discovery of novel compounds or repurposed drugs, which uses objectively measurable methodology [24]. This drug screening cascade performs drug assessments on metacestode vesicles (a and b), protoscoleces (c) and GL cells (d) [31–33]. These assays include drug activity assessments via different mechanisms (Fig 1). In (a) the activity of compounds against the metacestode stage is assessed by measuring the activity of the damage marker phosphoglucose isomerase (PGI), which is released into the supernatant of the cultured parasite upon impairment of metacestode vesicle integrity [33]. In addition, metacestode vesicle viability (b) upon treatment is assessed via measurement of ATP using CellTiter-glo assay [32]. In (c) the efficacy of compounds is measured via drug-induced reduction of protoscolex motility and morphological alterations [31]. The viability of isolated GL cells (d) upon treatment is also assessed via measurement of ATP using CellTiter-glo assay [32].

**Fig 1.**
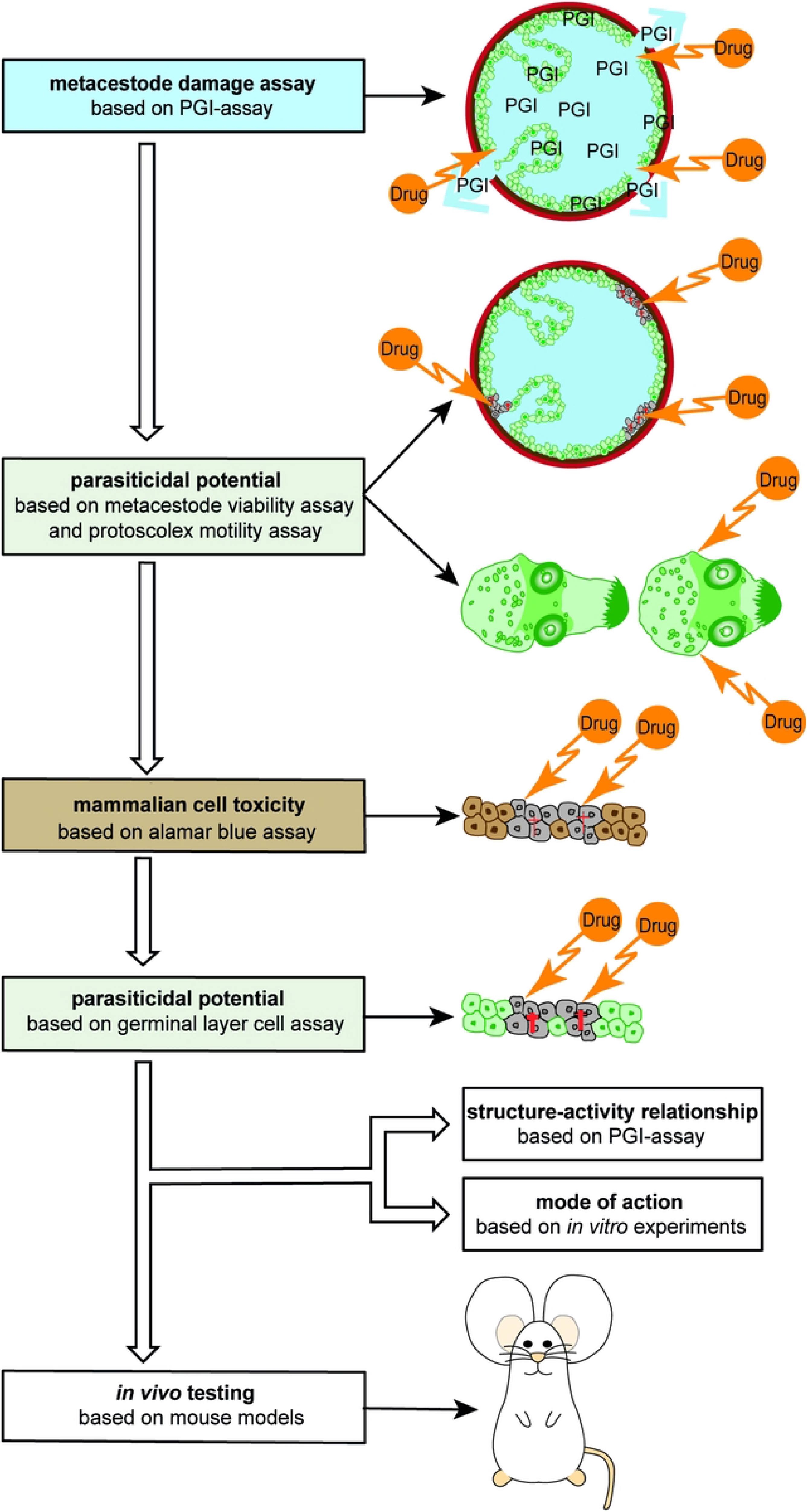
*In vitro* drug screening cascade for the identification of active compounds against *E. multilocularis* and *E. granulosus s.s.*. The drug screening cascade enables the identification of active compounds against *E. multilocularis* or *E. granulosus s.s.* metacestode vesicles, protoscoleces and GL cells via different *in vitro* assays. First, an overview screening is performed with metacestode vesicles to measure the direct impact on the physical integrity of metacestode vesicles via PGI assay [61], and a potential impairment of viability via vesicle viability assay [32]. Next, the damaging effect against protoscoleces is tested via motility assay [31], followed by cytotoxicity measurements in mammalian cells via alamar blue assays [61]. Finally, the parasiticidal potential of drug candidates against GL cells of the parasite are studied by measuring ATP levels [61]. Compounds that show parasiticidal activity against metacestode vesicles and GL cells, as well as a low toxicity against host cells, may be further studied for structure activity relationship via PGI assay and their mode of action via various specific *in vitro* assays [26,61]. Finally, promising compounds are tested in experimentally infected mice for their activity *in vivo* [32,63]. Figure adapted from [62].

In human patients, *E. granulosus s.l.* metacestode cysts grow between 1 and 50 mm in diameter a year and can be found in many different organs. However, in most cases either the liver or the lungs are affected [1,18]. The treatment of choice depends on the characteristics of the cysts, e.g. number, location and whether they are actively growing or not [18]. CE patients with inoperable cysts can be treated via chemotherapy with benzimidazoles (ABZ or mebendazole), or via PAIR (Puncture, Aspiration, Injection, Re-aspiration) [18,34]. However, even in cases where chemotherapy alone is not the preferred option, treatment with ABZ or mebendazole prior and after surgery or PAIR may facilitate the removal of the cyst and reduce the risk of secondary CE [34,35]. The current chemotherapeutic treatment of choice for inoperable cases is ABZ [18]. A review in which the response of CE patients to treatment with ABZ was evaluated found that 73.2 % of patients responded to treatment (either cured or improved), while 23.8 % showed either no change or a worsened condition [36]. These numbers demonstrate the need for new chemotherapeutic treatment options with better efficacy.

Previous *in vitro* drug screening methods for *E. granulosus s.l.* have been focusing on the demonstration of direct protoscolicidal effects mainly via the eosin exclusion test [37–40]. However, protoscoleces are not the disease-causing stage of the parasite, and the eosin exclusion test is based on subjective evaluation of eosin staining intensity by light microscopy. The morbidity of CE is caused by the growth of metacestode cysts that cause compression of affected organs [35]. Therefore, drug screening should primarily focus on the metacestode stage. While *in vitro* cultivation of *E. granulosus s.l.* metacestode cysts and subsequent drug testing have been described in the past, drug efficacy was always assessed via morphological readouts employing light microscopy and electron microscopy [41–43]. For the screening of higher numbers of compounds, electron microscopy is not feasible and light microscopy alone is prone to errors and misinterpretation. Thus, unbiased screening methods similar to the assays previously developed for *E. multilocularis* [31–33] should be made available.

While drug screening procedures should primarily target the metacestode vesicles as the disease-inflicting stage, it is important to consider GL cells in particular. The GL contains the stem cell population. The identification of compounds that kill stem cells will be crucial for future treatment options in order to decrease the risk of disease recurrence after treatment discontinuation [14]. Several attempts of isolating GL cells of *E. granulosus s.l.* metacestode vesicles or protoscoleces have been undertaken in the past [44–47]. However, isolating GL cells directly from hydatid cysts of infected hosts as it has been done in these studies bears a high risk of contamination with host cells [48]. Albani et al. (2010) reported on the culture of *E. granulosus s.l.* GL cells and these cultures were susceptible to 5-fluorouracil and paclitaxel, similar to protoscoleces and metacestode vesicles of *E. granulosus s.l.* [49]. 5-fluorouracil was later tested *in vivo* against *E. granulosus s.l.* in female CF-1 mice and showed activity that was similar to ABZ [50].

In this study, we have applied *in vitro* culture methods previously established for *E. multilocularis* to implement a standardized and reproducible protocol for the *in vitro* maintenance and propagation of *E. granulosus s.l.* metacestode vesicles and GL cells. In addition, we show that standardized and non-biased *in vitro* techniques for assessing the impact of drugs in *E. multilocularis* metacestode vesicles, GL cells and protoscoleces are also applicable for respective studies on *E. granulosus sensu stricto* (*s.s.*). parasite stages.

## Material and methods

### Chemicals and reagents

If not stated otherwise, all chemicals were purchased from Sigma-Aldrich (Buchs, Switzerland). Dulbeccos’s modified Eagle medium (DMEM) and Penicillin and Streptomycin (10’000 Units/mL Penicillin, 10’000 μg/mL Streptomycin) were purchased from Gibco (Fisher Scientific AG, Reinach, Switzerland). Fetal bovine serum (FBS) and Trypsin/EDTA (0.05 % Trypsin/ 0.02 % EDTA) were purchased from Bioswisstec (Schaffhausen, Switzerland). If not stated otherwise all plastic ware was purchased from Sarstedt (Sevelen, Switzerland). ABZ, buparvaquone (BPQ), mefloquine (MEF) and niclosamide (NIC) were all purchased from Sigma-Aldrich, MMV665807 (MMV-X) was from Princeton Biomolecular Research (South Brunswick Township, New Jersey, USA) and monepantel (MPT) was obtained from LuBioScience GmbH (Zürich, Switzerland). The drugs were prepared as 10.000 parts per million (ppm) and 40 mM stocks in DMSO and stored as aliquots at −20°C.

### Mice and ethics statement

For the maintenance of *E. multilocularis*, female BALB/c mice were purchased from Charles River Laboratories (Sulzheim, Germany) and used for experimentation after an acclimatization time of 2 weeks. The mice were maintained in a 12 h light/ dark cycle under a controlled temperature of 21 – 23°C, and a relative humidity of 45 – 55 % with food and water provided ad libitum. Each cage was enriched with a mouse house (Tecniplast, Gams, Switzerland), a tunnel (Zoonlab, Castrop-Rauxel, Germany) and nestlets (Plexx, Elst, Netherlands). All animals were treated in compliance with the Swiss Federal Protection of Animals Act (TSchV, SR455), and experiments were approved by the Animal Welfare Committee of the canton of Bern under the license numbers BE30/19 and BE2/2022.

#### Isolation of E. multilocularis and E. granulosus s.s. protoscoleces

Protoscoleces of *E. multilocularis* isolates MB17 and Smeen19 were isolated from metacestode cysts obtained from experimentally infected gerbils at the University of Würzburg, Germany. Protoscoleces of *E. granulosus s.s.* were aseptically aspirated from cysts obtained from livers or lungs of naturally infected sheep using a 50 mL syringe and an 18 g needle. The organs were sampled during routine veterinary *post mortem* examinations in two different abattoirs located in Sardinia, Italy: Buddusò (Sassari Province, 40.572532852821574, 9.25594927591384) and Lula (Nuoro Province, 40.395704769344924, 9.49113969820975). The sheep came from farms located in four different municipalities: Bonorva, Neoneli, Nulvi and Siniscola. In total, three different isolations were performed for *E. multilocularis* and four different isolations for *E. granulosus s.s.*.

#### Genotyping of *E. granulosus s.s.* isolates

All four *E. granulosus s.s.* isolations were genotyped using the mitochondrial targets cytochrome c oxidase I (*cox I*), NADH dehydrogenase I (*nad I*), the small ribosomal RNA *(rrnS*) and ATP synthase subunit 6 (*atp 6*) as previously described [51]. In short, sequences were amplified via PCRs with an initial heating step to 94°C for 3 minutes followed by 35 cycles of 94°C for 30 seconds, 56°C for 30 seconds and 72°C for 30 seconds. After those cycles, an additional elongation step at 72°C for 5 min was performed. Genotyping was performed for material of four isolations. The resulting sequences were aligned and cut to ensure same lengths and used to generate concatenated sequences with *atp 6* (328 bp) – *nad I* (514 bp) – *cox I* (337 bp) – *rrnL* (347 bp). The sequences were compared to concatenated sequences of the same lengths of *Taenia solium* [52], *E. multilocularis* [53], *E. oligarthrus* [54], *E. vogeli* [54] and *E. granulosus* genotypes G1 [55], G3 [56], G4 [55], G5 [54], G6 [54], G7 [54], G8 [54] and G10 [57]. A phylogenetic tree was generated with Molecular Evolutionary Genetics Analysis (MEGA) 11 [58] according to the maximum likelihood method using the Hasegawa-Kishino-Yano model (HKY) and gamma distributed with invariant sites (G+I). For the test of the phylogeny, the bootstrap method with 500 replications was chosen.

#### Culture of *E. multilocularis* metacestode vesicles

Metacestode vesicles of the isolate H95 were cultured as described by [27]. In short, metacestode material obtained from experimentally infected mice was pressed through a conventional tea strainer (Migros, Berne, Switzerland) and incubated overnight at 4°C in PBS containing penicillin (100 U/mL), streptomycin (100 µg/mL) and tetracycline (10 µg/mL). The next day, 1.5 mL of parasite material were co-cultured with semi confluent reuber rat hepatoma (RH) cells in DMEM containing 10 % FBS, penicillin (100 U/mL), streptomycin (100 µg/mL) and tetracycline (5 µg/mL). Once a week, the medium was changed and freshly trypsinized RH cells were added to the metacestode vesicles. In total, six independent metacestode vesicle cultures were generated.

#### Culture of *E. granulosus s.s.* metacestode vesicles

*E. granulosus s.s.* GL was aseptically removed from hydatid cysts from livers of naturally infected sheep. The GL was pressed through a conventional tea strainer (Migros, Berne, Switzerland) and incubated overnight at 4°C in PBS containing penicillin (100 U/mL), streptomycin (100 µg/mL), tetracycline (10 µg/mL) and an anti-contamination cocktail containing kanamycin (100 µg/mL), chloramphenicol (10 µg/mL) and flucytosine (50 µg/mL) [59]. The next day, 1.75 mL of parasite material were co-cultured with semi confluent RH cells in DMEM containing 10 % FBS, penicillin (100 U/mL), streptomycin (100 µg/mL), tetracycline (10 µg/mL) and an anti-contamination cocktail containing kanamycin (100 µg/mL), chloramphenicol (10 µg/mL) and flucytosine (50 µg/mL). Medium changes were performed as described for *E. multilocularis* and no additional anti-contamination cocktail was added after the first week. In total, two independent metacestode vesicle cultures were generated.

#### Isolation of GL cells of E. multilocularis and E. granulosus s.s

GL cells of *E. multilocularis* and, for the first time also for *E. granulosus s.s.,* were obtained from *in vitro* grown metacestode vesicles based on the protocol described by [28] with slight modifications. DMEM with 10 % FBS, penicillin (100 U/mL), streptomycin (100 µg/mL) and tetracycline (5 µg/mL) was conditioned by RH cells by incubation of 10^6^ cells in 50 ml medium for six days and 10^7^ cells in 50 mL medium for four days. These conditioned media were sterile filtered and mixed 1:1 (conditioned medium, cDMEM). Metacestode vesicles of either *E. multilocularis* or *E. granulosus s.s.* grown for at least one year were incubated in distilled water for two minutes to remove residual RH cells, subsequently washed in PBS and mechanically broken using a pipette, or a syringe and a conventional tea strainer, respectively, all at room temperature. The vesicle tissue was incubated in eight volumes of trypsin-EDTA solution at 37°C for 30 min and filtered through a 30 µm mesh (Sefar AG, Heiden, Switzerland). The remaining tissue was re-incubated in PBS until most of the cells were detached from the tissue which was visually confirmed by a reduced cloudiness of the PBS after each incubation step. Calcareous corpuscles were removed via centrifugation at 50 x g and the cell suspension was centrifuged at 600 x g for ten minutes. The pellet was taken up in cDMEM and the OD_600_ was measured of a 1:100 dilution. An OD_600_ value of 0.1 of the dilution corresponded to one arbitrary unit (AU) per µL of the undiluted cell suspension. 1000 AU of GL cells were incubated in five mL cDMEM per well of a 6-well plate at 37°C under a humid nitrogen atmosphere overnight. The next day, 2000 AU were united and let grown for three hours prior to preparation of the cells for the different experiments. Three different isolations were performed for each parasite species for the GL cell viability assay and one additional isolation for the vesicle formation assay.

#### Vesicle formation assay

GL cells of *E. granulosus s.s.* were incubated in 300 µL cDMEM containing 100 µM serotonin as this was shown to increase vesicle formation in *E. multilocularis* previously [60]. 150 AU of GL cells were seeded per well of a 96-well plate under a humid, microaerobic atmosphere (85 % N_2_, 10 % CO_2_, 5 % O_2_) and cultured for 26 days. Three times a week, half of the medium was changed and photos were taken to follow vesicle formation of primary cell cultures after five, twelve and 26 days on a Nikon TE2000E microscope connected to a Hamatsu ORCA ER camera.

### Transmission electron microscopy

Metacestode vesicles of *E. granulosus s.s.* were grown for 4 months *in vitro* and then processed for transmission electron microscopy as previously described with few modifications [61]. The samples were fixed in 100 mM sodium-cacodylate (pH 7.3) containing 2 % glutaraldehyde at 4°C overnight. After three washing steps 100 mM sodium-cacodylate (pH 7.3), the samples were post-fixed in 100 mM sodium-cacodylate (pH 7.3) containing 2 % osmium tetroxide at RT for 1.5 hours. Then, the samples were washed in water and dehydrated stepwise using a series of washing steps in ethanol (30 %, 50 %, 70 %, 90 %, three times 100 %). The samples were embedded in Epon 812 resin and incubated at 37°C with two subsequent resin changes in a total incubation time of 2.5 hours. Next, the samples were left in resin for 24 hours at RT and then incubated to 65°C overnight for polymerization. Finally, 80 nm sections were cut with an ultramicrotome (Reichert and Jung, Vienna, Austria), and were loaded onto formvar-carbon coated nickel grids (Plano GmbH, Marburg, Germany). The samples were stained with Uranyless™ and lead citrate (Electron Microscopy Sciences, Hatfield PA, USA) and were photographed on a FEI Morgagni transmission electron microscope (Field Electron and Ion Company, Hillsboro, Oregon, USA) operating at 80 kV.

### In vitro drug screening cascade for E. multilocularis and E. granulosus s.s

#### Overview of the *in vitro* drug screening cascade for *E. multilocularis* and *E. granulosus s.s*

In order to identify novel compounds with activity against *Echinococcus*, the *in vitro* drug screening cascade previously developed for assessing drugs against metacestode vesicles and stem cells of *E. multilocularis* was further refined and applied to *E. granulosus s.s.* in this study [24,62]. First, the effects of compounds on the physical integrity of the disease-causing metacestode vesicles was measured via phosphoglucose isomerase (PGI) assay) [61], followed by measuring the effects of compounds on the viability of the metacestode vesicle tissue [32]. Upon cyst rupture during *E. granulosus s.l.* infection, protoscoleces in human patients can cause the formation of secondary cysts [35], thus we added the protoscolex motility assay [31] to identify compounds that also show activity against this stage (Fig 1). Active compounds can then be further assessed for cytotoxicity or viability impairment in mammalian cells, and against isolated GL cells of the parasite [61]. If the different assays propose a therapeutic window, with high anti-parasitic activity and low toxicity against mammalian cells, promising compounds may be tested in the murine models of primary and secondary echinococcosis, reflecting early and late infection stages [32,63].

### Phosphoglucose isomerase assay

The PGI assay allows the assessment of drug-induced damage on the integrity of metacestode vesicles of *E. multilocularis,* which can be measured indirectly via the leakage of vesicle fluid containing EmPGI into the culture medium. PGI assays were performed as previously described [61] with the exception that metacestode vesicles were incubated in a microaerobic atmosphere (85 % N_2_, 10 % CO_2_, 5 % O_2_) instead of a normoxic atmosphere (5% CO_2_). This reflects closer the situation these parasites encounter *in vivo* [64]. In short, metacestode vesicles of around two mm in diameter were purified via several washing steps in PBS and mixed with two volumes of DMEM without phenol red containing penicillin (100 U/mL) and streptomycin (100 µg/mL). The metacestode vesicles were distributed to a 48-well plate (Huberlab, Aesch, Switzerland) with one mL per well. Based on previous studies [31,61,63,65], drugs were added in triplicates in 0.1 % DMSO to a final concentration of 40 µM, except MMV-X and NIC, which were added to a final concentration of 1 µM due to their higher reported activity against *E. multilocularis* metacestode vesicles [62]. 0.1 % TX-100 was used as a positive control. Measurements were performed on an EnSpire multilabel reader (Perkin Elmer, Waltham, MA, USA). Three independent experiments from three independent metacestode vesicle batches for each parasite species were performed, and mean values and SDs are given. In PGI assays, compound concentrations were regarded as active against metacestode vesicles when inducing a minimum of 20 % PGI release compared to the positive control. Representative photographs of the metacestode vesicles were taken after five and twelve days using a Nikon SMZ18 stereo microscope (Nikon, Basel, Switzerland) at 0.75X magnification. Drug activity between *E. multilocularis* and *E. granulosus s.s.* metacestode vesicles was compared by multiple two-tailed students t-tests with equal variances using mean values of three independent experiments. Bonferroni-corrected *p* values of *p<*0.05 were considered significant for all tests.

### Metacestode vesicle viability assay

This assay was done to investigate whether a drug was not only able to rupture the integrity of the metacestode vesicle, but also to impair the viability of the GL layer tissue inside the metacestode vesicle. Metacestode vesicle viability was assessed in the same plates used for the PGI assay after 12 days of incubation. TX-100 at a final concentration of 0.1 % was added to all wells and metacestode vesicles were mechanically broken using a pipette. 50 µL of supernatant were mixed with 50 µL of CellTiter-Glo (Promega, Dübendorf, Switzerland) and measurements were performed on an EnSpire multilabel reader. Values are shown in relation to the DMSO control and mean values and SDs of three independent experiments from three independent metacestode vesicle batches for both parasite species are shown. For metacestode vesicle viability assessments, compounds were identified as active when causing at least a reduction of 50 % viability in relation to the DMSO control treatment. Metacestode vesicle viability of *E. multilocularis* and *E. granulosus s.s.* metacestode vesicles was compared by multiple two-tailed students t-tests with equal variances using mean values of three independent experiments. Bonferroni-corrected *p* values of *p<*0.05 were considered significant for all tests.

### Protoscolex motility assay

Protoscolex motility assay allows the objective assessment of drug-induced motility reduction of protoscoleces when incubated with different compounds. It was performed as previously described [31]. In short, protoscoleces of *E. multilocularis* were obtained from metacestodes isolated from experimentally infected gerbils, and *E. granulosus s.s.* protoscoleces were isolated from metacestodes collected from naturally infected sheep at abattoirs in Sardinia. The protoscoleces were separated from the GL via filtering through a 250 µm mesh, followed by several washing steps in PBS. Then the protoscoleces were activated with 10 % DMSO at 37°C for three hours. Activated protoscoleces were incubated in DMEM containing 10 % FBS, penicillin (100 U/mL) and streptomycin (100 µg/mL) overnight at 37°C under 5 % CO_2_ atmosphere. The next day, protoscoleces were washed in PBS and mixed with DMEM without phenolred containing 10 % FBS and distributed in a 384 well plate (Huberlab, Aesch, Switzerland) with compounds at a final concentration of 100 to 0.0006 ppm for praziquantel (PZQ) and its enantiomers or 100 to 0.04 ppm for the other drugs, respectively (six replica per concentration) with 1 % DMSO each, and in a final volume of 20 µL. The concentration of the drugs was chosen to be given in ppm to have a direct comparison to previous published results [31]. The plates were sealed with a clear view seal foil (Huberlab, Aesch, Switzerland) and incubated at 37°C. After 12 hours, photographs were taken with ten second intervals using a Nikon TE2000E microscope connected to a Hamatsu ORCA ER camera at 40 times magnification. The software NIS Elements Version 4.40 with the additional module JOBS was used. The motility of protoscoleces was assessed by subtracting the images from each other via an automated script in imageJ [31]. The motility was shown in percentage of the DMSO control (1 %) as mean values of three independent experiments (including protoscoleces of independent isolations) with standard deviations (SD). Those compounds that caused an impairment of motility by 50 % in relation to the DMSO control were designated as active. Concentration-dependent reduction of the motility of *E. multilocularis* and *E. granulosus s.s.* protoscoleces was compared by multiple two-tailed students t-tests with equal variances using mean values of three independent experiments. One test was performed per concentration of a drug and thus the *p* values were bonferroni corrected. Bonferroni-corrected *p* values of *p<*0.05 were considered significant for all tests.

### GL cell viability assay

Isolated *E. multilocularis* and *E. granulosus s.s.* GL cells were distributed into wells of black 384-well plates as 15 A.U. per well in 12.5 µL cDMEM. Compounds were added to a final concentration of 40 µM for all drugs, expect MMV-X and NIC, which were tested at 1 µM in quadruplicates in another 12.5 µL cDMEM. 0.1 % DMSO was used as a negative control. *Echinococcus* cells were incubated under a humid, microaerobic atmosphere for five days and pictures were taken at the end of the incubation on a Nikon TE2000E microscope connected to a Hamatsu ORCA ER camera at 40 times magnification. 25 µL of CellTiter-Glo (Promega, Dübendorf, Switzerland) containing 1 % TX-100 was added to the wells and cell aggregates were disrupted by pipetting. Measurements were performed on a HIDEX Sense microplate reader (Hidex, Turku, Finland). Values are shown in relation to the DMSO control and mean values and SDs of three independent experiments from three independent GL cell isolations for both parasite species are shown. Efficacy of drugs between *E. multilocularis* and *E. granulosus s.s.* GL cells was compared by performing multiple two-tailed students t-tests with equal variances with mean values of three independent experiments. The calculated *p* values were bonferroni-corrected. Bonferroni-corrected *p* values of *p<*0.05 were considered significant for all tests.

## Results

### Genotyping of *E. granulosus s.s.* isolates

Genotyping was performed for all four isolates of *E. granulosus s.s.* and showed that the genotypes of three isolates were G1 and one isolate was G3 (Supplementary Fig 1).

#### Dedifferentiation of *E. granulosus s.s.* protoscoleces into metacestode vesicles

In order to generate *E. granulosus s.s.* metacestode vesicles in sufficient numbers for subsequent drug screening assays, we cultured GL of *in vivo* grown *E. granulosus s.s.* hydatid cysts with RH cells as published for *E. multilocularis*. After three weeks, protoscoleces had de-differentiated into metacestodes vesicles (Fig 2A) and after five weeks those vesicles had developed a LL clearly visible by light microscopy (Fig 2B). Metacestode vesicle cultures were cultured *in vitro* for prolonged times and respective images are shown after ten months in culture (Fig 2C) and twenty months in culture (Fig 2D). The presence of the LL was confirmed by TEM after four months in culture (Figs 2E and 2F).

**Fig 2.**
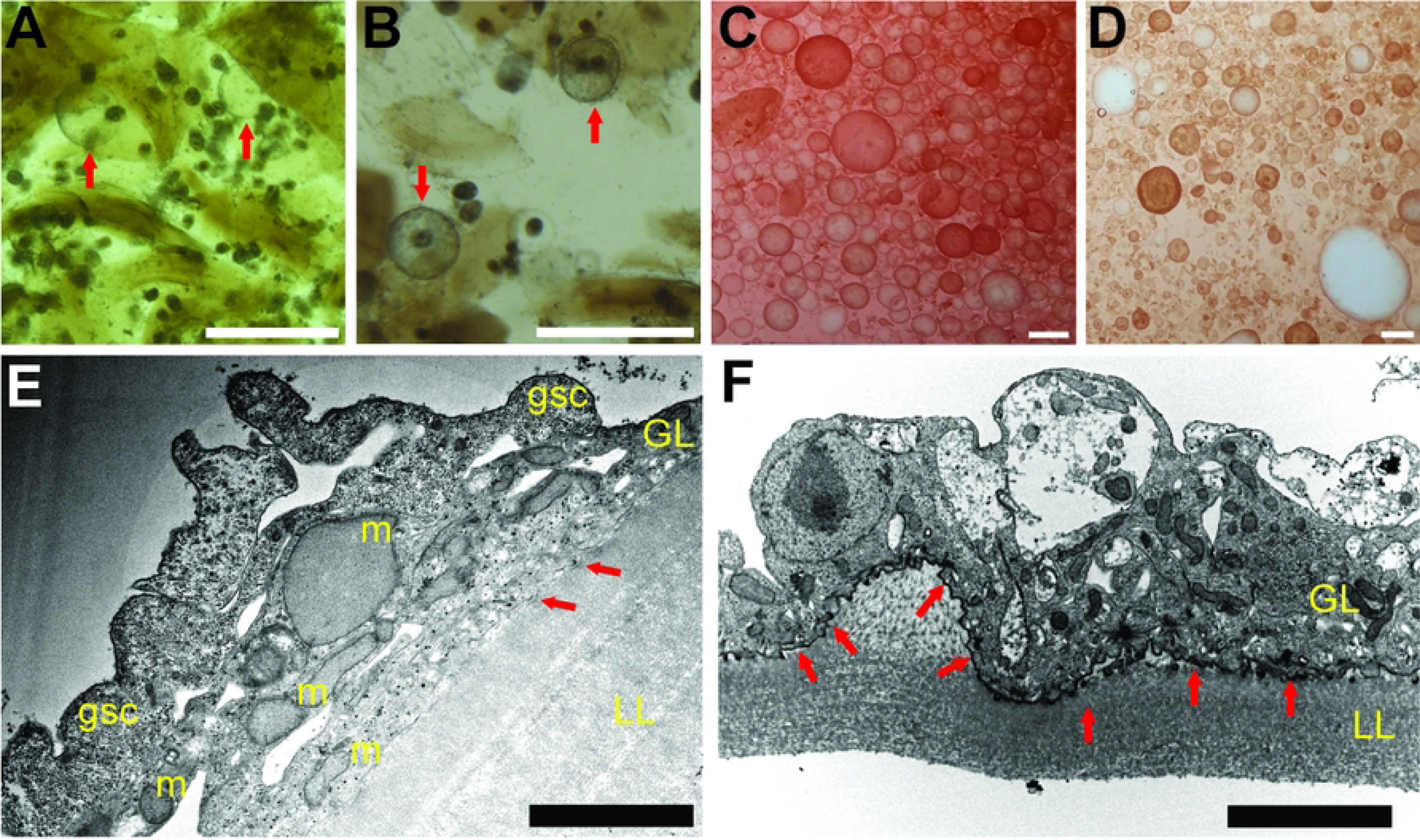
*In vitro* cultured *E. granulosus s.s.* metacestode vesicles. *E. granulosus s.s.* metacestode vesicles were generated upon *in vitro* culture of GL from *ex vivo* hydatid cysts. A: Metacestode vesicles appeared after three weeks as indicated by arrows. B: After five weeks, vesicles generated a visible laminated layer as indicated by arrows. C: Metacestode vesicles after ten months of culture. D: Metacestode vesicles after 20 months of culture. Scale bars represent one mm in A and B or five mm in C and D. E and F: TEM of *E. granulosus s.s.* metacestode vesicles after four months of culture. LL = laminated layer; GL = germinal layer; gsc = glycogen storage cell; m = mitochondrion, nuc = nucleus. Note the presence of very short microtriches (arrows), and the absence of a discernible tegument at the LL-GL interface. Scale bars represent 4 µm and 8 µm in E and F, respectively.

#### Drug efficacy against metacestode vesicles measured by PGI assay

The efficacy of a set of standard drugs was assessed using *in vitro* cultured *E. multilocularis* and *E. granulosus s.s.* metacestode vesicles. The conditions previously described for *E. multilocularis* were applied to *E. granulosus s.s.* metacestode vesicles, with the positive control TX-100 causing maximum physical damage. Active drugs led to rupture of metacestode vesicles after five days of culture, whereas metacestode vesicles treated with DMSO stayed intact during at least twelve days (Figs 3A and 3B). Drug-induced PGI release was similar in both species upon treatments with ABZ, MEF, NIC and NTZ after five and twelve days of drug incubation (Figs 3C and 3D). MMV-X displayed higher activity against *E. multilocularis* metacestode vesicles after five days (*p*=0.001), but this difference was smaller after twelve days. In contrast, MPT exerted a similar effect in both parasites after five days, but exhibited a significantly higher activity against *E. multilocularis* metacestode vesicles after twelve days (*p*=0.039). BPQ exerted slightly higher activity against *E. multilocularis* metacestode vesicles after five and twelve days, but differences were not significant.

**Fig 3.**
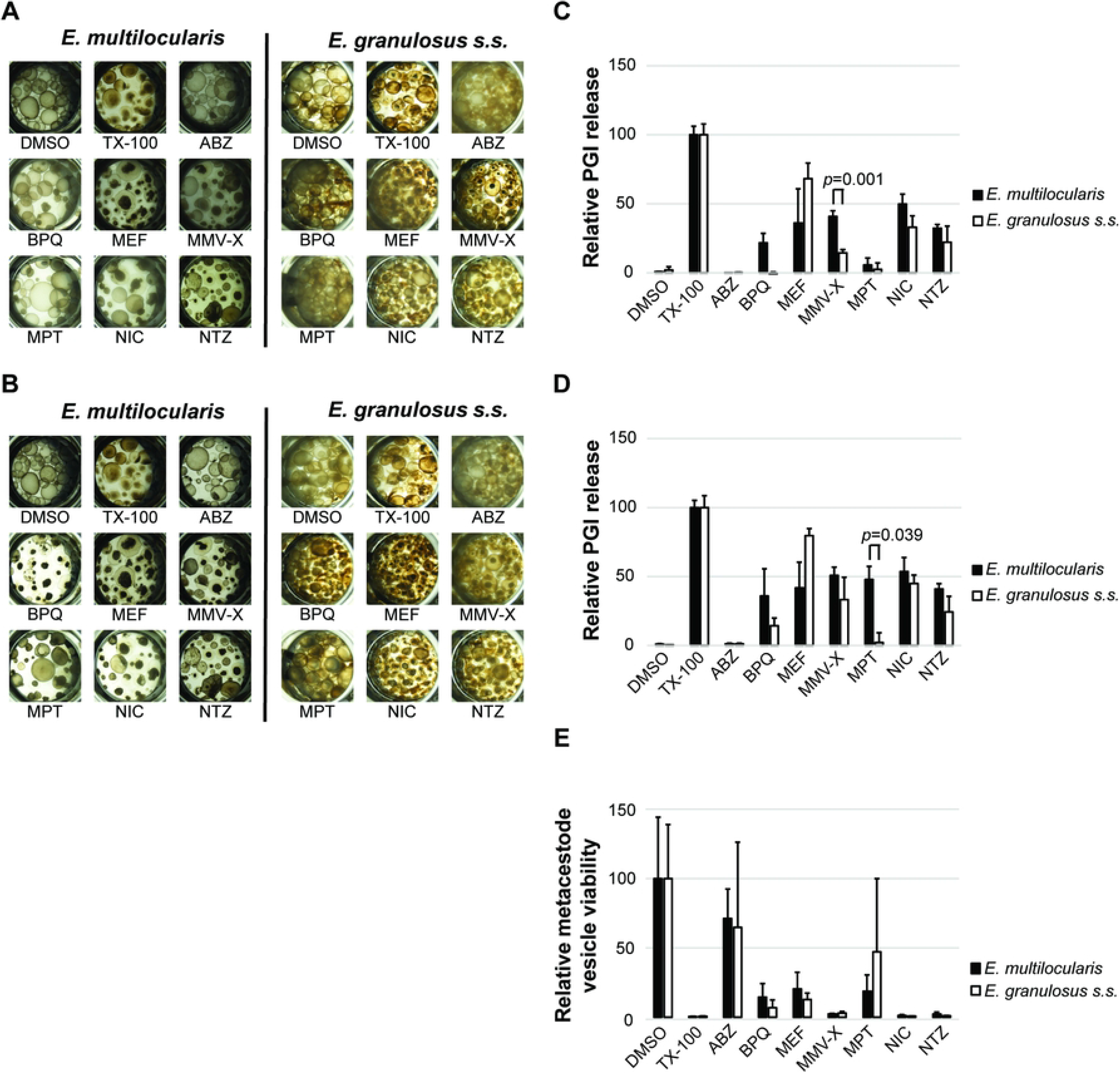
Assessment of drug efficacy on *E. multilocularis* and *E. granulosus s.s.* metacestode vesicles by PGI assay and metacestode vesicle viability assay. PGI release of *E. multilocularis* and *E. granulosus s.s.* metacestode vesicles into culture supernatant was measured after five days (C) and twelve days (D) and is shown in relation to the positive control 0.1% TX-100. Representative images of *E. multilocularis* and *E. granulosus s.s.* metacestode vesicles treated with the negative control 0.1 % DMSO, the positive control 0.1 % TX-100, or different drugs after five days (A) and twelve days (B) are shown. Metacestode vesicle viability was measured after twelve days and is shown in relation to the negative control 0.1% DMSO (E). The activities of ABZ (albendazole), BPQ (buparvaquone), MEF (mefloquine), MMV-X (MMV665807), MPT (monepantel), NIC (niclosamide) and NTZ (nitazoxanide) were assessed *in vitro* against metacestode vesicles of *E. multilocularis* and *E. granulosus s.s.*. Drug activity was calculated in percentage as relative PGI release compared to the positive control 0.1 % TX-100. The drugs were tested at 40 µM concentrations, except MMV-X and NIC, which were tested at 1 µM. The metacestode vesicles were incubated for five and twelve days under a humid microaerobic atmosphere (85 % N_2_, 10 % CO_2_, 5 % O_2_) and tests were performed in triplicates per condition. Shown are mean values and standard deviations of three independent experiments. Relative PGI release of *E. multilocularis* and *E. granulosus s.s.* metacestode vesicles was compared and significance is shown according to bonferroni-corrected *p* values of *p* <0.05 obtained using multiple two-tailed students t-tests assuming equal variance.

#### Drug efficacy against metacestode vesicles measured by viability assay

The metacestode vesicle viability assay previously developed for *E. multilocularis* was also transferable to *E. granulosus s.s.*. DMSO had no impact, while TX-100 and active drugs impaired viability (Fig 3E). The active drugs BPQ, MEF, MMV-X, NIC and NTZ caused a strong reduction of metacestode vesicle viability in both species, whereas ABZ had no impact. Interestingly, exposure of metacestode vesicles to MPT displayed no measurable effect in the PGI assay, but led to reduced metacestode vesicle viability upon measurement of ATP using CellTiter-glo assay. No significant differences for the drug activities were detected between *E. multilocularis* and *E. granulosus s.s.* in this assay.

#### Drug efficacy against protoscoleces measured by motility assay

Protoscolex motility assays were carried out with protoscoleces of both species. They were incubated with various drugs published to be active against *E. multilocularis* protoscoleces (NIC, NTZ and MMV-X) or known to have no activity (ABZ and MPT) (Fig 4). Dose-response curves showed no significant differences for MMV-X, MPT, NIC and NTZ between *E. multilocularis* and *E. granulosus s.s.* protoscoleces. At a very high concentration of 100 ppm (377 µM), ABZ was significantly more active against *E. granulosus s.s.* protoscoleces (*p*=0.001).

**Fig 4.**
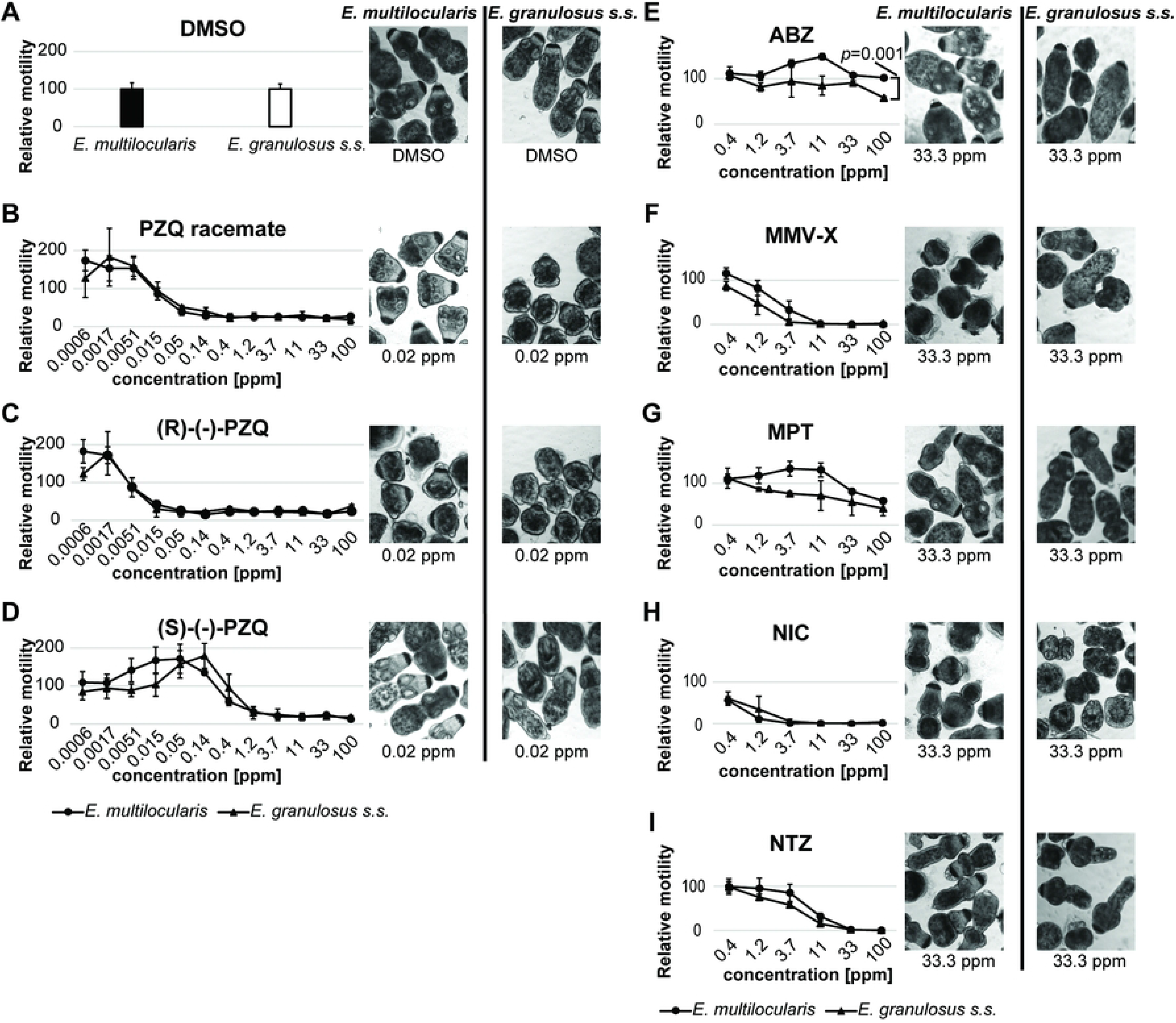
Effects of drugs on *E. multilocularis* and *E. granulosus s.s.* protoscolex motility. The activities of PZQ (praziquantel) racemate (B), as well as of (R)-(-)-PZQ (C), (S)-(-)-PZQ (D) and various standard drugs (ABZ (albendazole) (E), MMV-X (MMV665807) (F), MPT (monepantel) (G), NIC (niclosamide) (H), and NTZ (nitazoxanide) (I)) were assessed. The relative motility is shown in comparison to the solvent control (1 % DMSO). The morphology of protoscoleces treated with PZQ and its enantiomers is shown for 0.02 ppm and for protoscoleces treated with the other drugs, the morphology is shown for 33.3 ppm. All concentrations were tested in six replica. Shown are mean values and standard deviations of three independent experiments. Significant bonferroni-corrected *p* values with *p*<0.05 are shown that were obtained using students t-test comparing relative movement between *E. multilocularis* and *E. granulosus s.s.* protoscoleces for each individual drug concentration.

Additionally, a racemic mixture of PZQ, as well as of its R-enantiomer and S-enantiomer, were assessed. Protoscoleces of both species showed very similar dose-dependent response curves to these drugs (Fig 4). As previously published for *E. multilocularis* [31], none of these drugs completely inhibited the motility of *E. granulosus s.s.* protoscoleces. However, (S)-(-)-PZQ was largely inactive, whereas (R)-(-)-PZQ was the active form. The activity of these compounds was also visible upon morphological assessment of protoscoleces. Alterations were observed in both species for PZQ racemate and (R)-(-)-PZQ, but not for (S)-(-)-PZQ at 0.02 ppm (0.05 µM) after 12 hours of incubation (Fig 4). No significant differences were found between *E. multilocularis* and *E. granulosus s.s.* protoscoleces.

### Culture and drug efficacy measurements employing GL cells

#### *E. granulosus s.s.* primary cells have the capacity to form new metacestode vesicles

In order to determine whether the here described protocol of GL cell isolation for *E. granulosus s.s.* would result in viable cell cultures, respective GL cells were maintained *in vitro* for up to 26 days. After 5 days, microscopy revealed the appearance of vesicular structures reminiscent of metacestode vesicles, which could be clearly distinguished from the surrounding aggregates after twelve days (Fig 5). These newly formed *E. granulosus s.s.* metacestode vesicles did not rapidly increase in size but separated more clearly from the aggregates after 26 days of *in vitro* culture.

**Fig 5.**
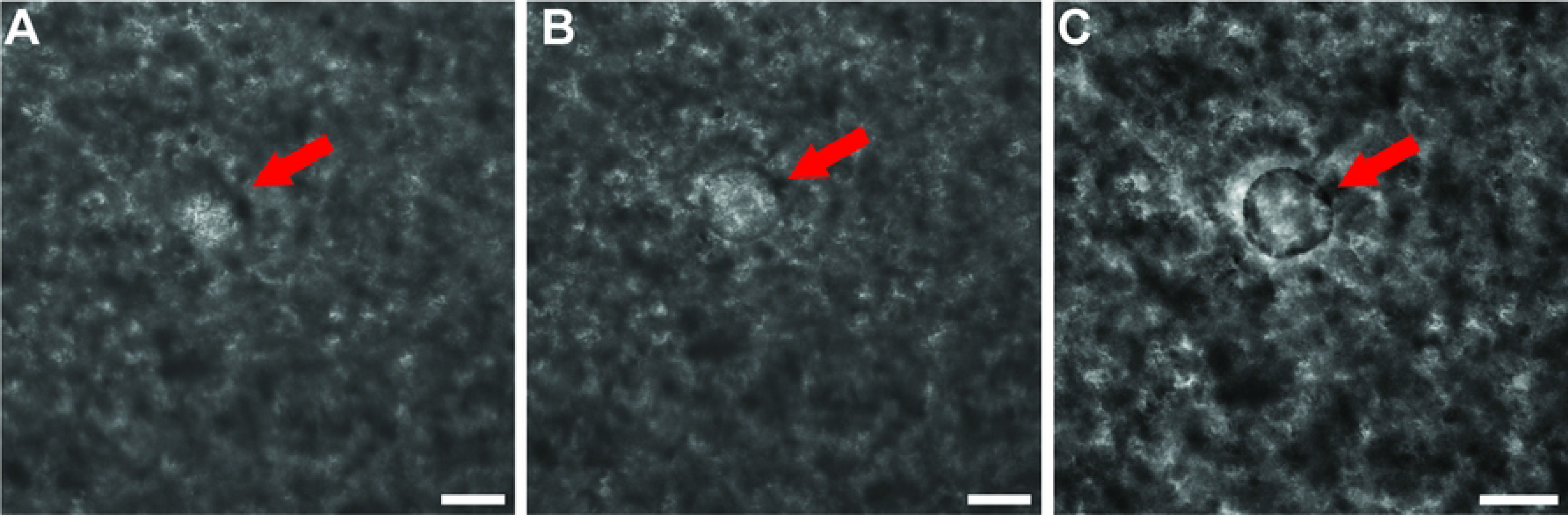
Metacestode vesicle formation from *E. granulosus s.s.* GL cells. *E. granulosus s.s.* GL cells were cultured in cDMEM supplemented with 100 µM serotonin with three medium changes a week under microaerobic conditions. Micrographs were taken after five days (A), twelve days (B) and 26 days (C). Scale bars represent 100 µm. Red arrows highlight the position of the metacestode vesicle.

#### Drug efficacy assessment by GL cell viability assay

The same panel of seven standard drugs was tested against isolated GL cells of both parasites (Fig 6). In the DMSO control these GL cell cultures formed aggregates, but in the presence of active drugs, this process was inhibited. Very similar viability impairment was measured when *E. multilocularis* and *E. granulosus s.s.* GL cells were cultured in the presence of BPQ, MEF, MMV-X and NIC. Interestingly, ABZ and NTZ had a significantly higher activity against *E. multilocularis* GL cells in this *in vitro* setup within five days of incubation (*p*=0.0002 and *p*=0.002, respectively). MPT was slightly more active against *E. multilocularis* GL cells, but this difference was not significant.

**Fig 6.**
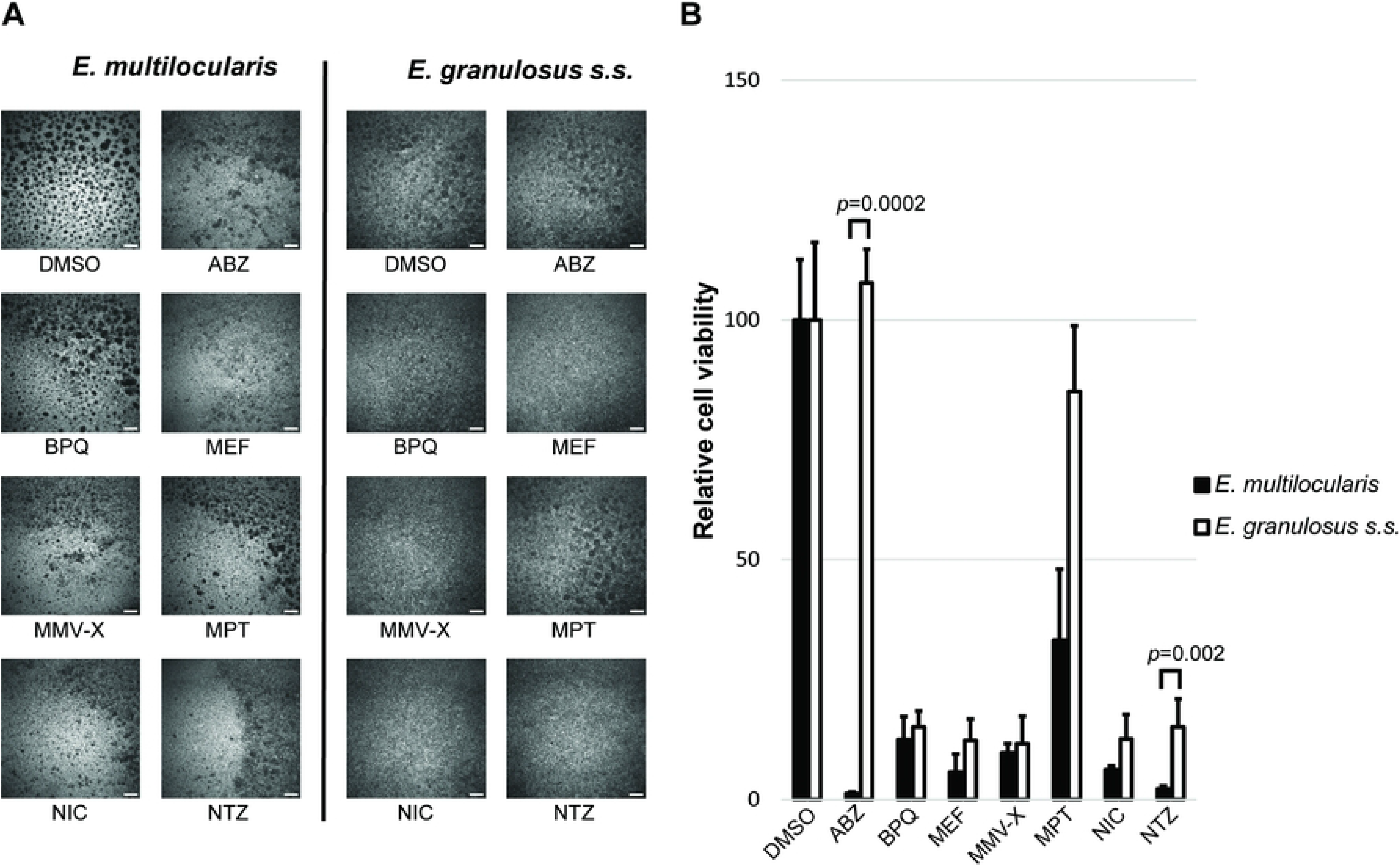
Efficacy of drugs against *E. multilocularis* and *E. granulosus s.s.* GL cells. The activities of ABZ (albendazole), BPQ (buparvaquone), MEF (mefloquine), MMV-X (MMV665807), MPT (monepantel), NIC (niclosamide) and NTZ (nitazoxanide) were assessed for *E. multilocularis* and *E. granulosus s.s.* primary cells. The relative cell viability after five days of incubation with the compounds is shown in comparison to the solvent control (0.1 % DMSO) (B) and representative images are shown with scale bars representing 200 µm (A). Drugs were tested at 40 µM concentrations, except MMV-X and NIC, which were tested at 1 µM. For each experiment, 15 AU of primary cells of either *E. multilocularis* or *E. granulosus s.s.* were incubated in cDMEM containing the respective compounds. The plate was incubated for 5 days under a humid microaerobic atmosphere (85 % N2, 10 % CO2, 5 % O2) and performed in four replica per condition. Shown are mean values of three independent experiments. Significant bonferroni-corrected *P* values with *p*<0.05 are shown that were obtained using multiple students t-test comparing relative cell viability between *E. multilocularis* and *E. granulosus s.s.* GL cells for the different drugs.

## Discussion

The neglected tropical diseases AE and CE are caused by the tapeworms *E. multilocularis* and *E. granulosus s.l.*. Respective metacestodes inflict serious clinical manifestations, and new parasitocidal drugs are needed for curative treatment. The search for new drugs can focus on repurposing of marketed drugs or on evaluating the efficacy of novel compounds against these parasites [24]. In any case, drugs have to show activity in whole-organism based systems against the different larval stages and specifically against the stem cells of *Echinococcus*. It has been speculated that stem cells within metacestodes are unaffected by the currently used parasitostatic benzimidazoles and thus are likely to decisively contribute to the observed recurrence of the disease after treatment discontinuation [22,24].

For *E. multilocularis,* an in *in vitro* drug screening cascade was established previously, which allows to assess the efficacy of drugs of interest against metacestodes, protoscoleces and GL cells in an unbiased manner, and includes the evaluation of the parasitocidal potential of compounds [62]. It has to be noted that these screening assays show the direct effect of a compound against different stages of parasites *in vitro,* but potential indirect effects that require previous metabolization of a compound, or the functional immune system of a host, are missed [66].

Despite the much higher numbers of human CE cases compared to AE, a similar drug screening cascade has so far not been applied to *E. granulosus s.l.*. This paper closes this gap. The here applied *E. granulosus s.s.* isolates from Sardinia were of the genotypes G1 and G3 which are the most common genotypes found in Sardinia [67,68]. G1 causes 88.4 % of human CE cases worldwide [69]. However, we speculate that the very same assays should also be applicable to genotypes G6 and G7, which are responsible for 7.3 % and 3.7 % of human infections worldwide, respectively [69]. Future evaluation of our assays with G1 and G3 from other regions and different animal origin, as well as screening assays with G6 and G7 will show whether the methodology used herein can be applied to all *E. granulosus s.l.* genotypes.

Medium throughput screening of anti-echinococcal compounds requires large amounts of metacestode vesicles [24]. For *E. multilocularis*, optimized culture systems, in which homogenized parasite material is co-cultured with RH cells, have allowed the large-scale generation of metacestode vesicles *in vitro* [28]. For *E. granulosus s.l.*, the *in vitro* generation of metacestode vesicles is described using activated protoscoleces without feeder cells [42,43,70], or with murine Hepa 1-6 cells [71]. In our study, we reliably generated large amounts of *E. granulosus s.s.* metacestode vesicles starting from homogenized GL that could be cultivated for at least 20 months together with RH cells.

For *E. multilocularis,* previous studies showed that activity of the drugs MMV-X, MEF and a series of endochin-like quinolones was similar for GL cells and metacestode vesicles [32,72]. For *E. granulosus s.l.*, similar observations were reported using cells directly isolated from hydatid cysts grown in cattle and treated with the drugs 5-fluorouracil and paclitaxel [49]. In order to reduce the risk of co-isolating host cells, we here isolated GL cells of *E. granulosus s.s.* metacestode vesicles that were grown for at least one year *in vitro* and that were treated with distilled water to kill any RH cells potentially still attached to the parasites. To the best of our knowledge, this is the first report of *E. granulosus s.s.* GL cells isolated from *in vitro* generated metacestode vesicles and the use of these cells for long-term culture and drug testing. The respective GL cells formed aggregates and developed new metacestodes vesicles *in vitro* similar to the situation previously reported for *E. multilocularis* [25,26]. So far, an appearance of cystic-like structures from *E. granulosus s.s.* cells has only been described from a cell line cultured in biphasic medium [47]. Here, we present the formation of metacestode vesicles from GL cell cultures using cells isolated from *in vitro* generated metacestode vesicles instead of from *in vivo* grown hydatid cysts.

By applying the PGI assay and the metacestode vesicle viability assay, the standard drugs BPQ, MEF, MMV-X, NIC and NTZ exhibited distinct activities under microaerobic conditions. This is in accordance to previous studies, where these drugs were published to show activity against *E. multilocularis* metacestode vesicles under aerobic or anaerobic conditions [61–63,65]. We subjected isolated GL cells of *E. granulosus s.s.* and *E. multilocularis* to viability assays in order to assess whether their response to drugs was in line with our results from the PGI assay and the metacestode vesicle viability assay. We showed that MEF, NIC and NTZ were also active against isolated *E. granulosus s.s.* GL cells. This was also the case for BPQ and MMV-X, however, clear activity in the PGI assay against *E. granulosus s.s.* metacestode vesicles was apparent only after 12 days. MPT was not active against *E. granulosus s.s.* metacestode vesicles in the PGI assay, but showed activity in the metacestode vesicle viability assay. In contrast, no activity could be detected when assessed in isolated GL cell cultures. This could possibly be explained by the differences in incubation times for these viability assays, with twelve days of drug treatment for metacestode vesicles compared to five days for GL cell cultures. MPT had a stronger effect against *E. multilocularis* as determined by PGI and metacestode vesicle viability assay after 12 days, and in GL cell cultures as well.

The current drug of choice for the treatment of AE and CE patients, ABZ, was not active against *E. multilocularis* metacestode vesicles *in vitro* in our assay system. While this might come as a surprise, it confirms previous studies showing that under *in vitro* conditions longer incubation times are needed to assess benzimidazole-activity against metacestode vesicles [33,73]. In fact, previous reports noted that ABZ and fenbendazole treatments induce the loss of microtriches at the tegument-laminated layer interface, and thus reduce the absorbing surface area of the parasite. While microtriches were shortened or largely lost as an immediate drug-response, the physical integrity of the residual tegument remained intact, and PGI-release was not measurable until later timepoints [74,75]. Interestingly, we here observed activity of ABZ against GL cells of *E. multilocularis*, a finding that was neither reported, nor assessed in previous studies. In contrast, ABZ did not exhibit any effects when applied for the treatment of *E. granulosus s.s.* metacestode vesicles or GL cells under the conditions used. A possible explanation could be that *E. granulosus s.s.* GL cells possibly exhibit a slower proliferation rate. Future projects will have to assess the efficacy and modes of action of benzimidazoles against *Echinococcus in vitro* in more depth.

The activity of MMV-X, NIC and NTZ against protoscoleces of *E. granulosus s.s.* and *E. multilocularis* were in line with activity against metacestode vesicles or GL cells. As expected, ABZ showed no activity against *E. granulosus s.s.* and *E. multilocularis* protoscoleces, and MPT was active only against *E. granulosus s.s.* protoscoleces at 100 ppm (211 µM), which is a rather high concentration. BPQ and MEF were not tested against protoscoleces as they were not part of the previous drug screening against *E. multilocularis* protoscoleces that served as an internal control [31]. Concentration-dependent activities for *E. multilocularis* protoscoleces were very similar to those obtained in previous studies. Slight differences were observed for PZQ racemate that showed activity at 0.05 ppm (0.15 µM) instead of 0.02 ppm (0.05 µM), for ABZ that showed no activity at all against *E. multilocularis* protoscoleces compared to little activity at 100 ppm (377 µM) in previous results and for NTZ that already showed activity at 11.1 ppm (36 µM) instead of at (108 µM).

The inter-species comparison of the activities of the different drugs shows that ABZ and NTZ exhibited higher activities when tested in *E. multilocularis* GL cell cultures, and MMV-X and MPT displayed higher activity against *E. multilocularis* metacestode vesicles when compared to *E. granulosus s.s.*. ABZ was not active against *E. multilocularis* protoscoleces, but *E. granulosus s.s.* protoscoleces were susceptible to ABZ, albeit at a very high concentration. PZQ and its enantiomers displayed similar concentration-dependent activities in both *Echinococcus* species.

Overall, we here show that the *in vitro* drug screening cascade previously established for the assessment of interesting compounds against *E. multilocularis* [31–33] can be applied to *E. granulosus s.s.* metacestode vesicles, GL cells and protoscoleces, and serves as a whole-organism based tool to test anti-echinococcal activity of novel compounds. As an additional early step, the toxicity against mammalian cells must be analyzed to test for a potential therapeutic window. Toxicity assays employing mammalian cells, were not included in this work as this has been done before for MMV-X [32] and all other compounds are marketed drugs considered safe for human or animal use. Once a therapeutic window is established, meaning that compounds display high anti-parasitic activity and comparatively minimal cytotoxicity, one can prospectively proceed to assessment in the murine models of AE [76–81]. Whether compounds showing profound activity against *E. granulosus s.s.* in this *in vitro* screening cascade are also active *in vivo* will be investigated in the future applying established CE mouse models [82–84].

In conclusion, we have shown that the application of robust assays published for protoscoleces, metacestode vesicles and GL cells of *E. multilocularis* [31,33,32] can be applied to *E. granulosus s.s.*, which allows the comparative analysis of drug activities between the two *Echinococcus* species. The culture methods described herein allow the generation of large amounts of *E. granulosus s.s.* metacestode vesicles and pure, proliferative GL cells which is a prerequisite for medium throughput drug screening assays. Future drug screening projects should include all these stages of *E. granulosus s.s.*, because metacestodes are the disease-causing stage, but protoscoleces can develop into secondary cysts when spilled either spontaneously or during surgical removal of cysts [35]. It was shown that combined treatment of ABZ and PZQ is superior in reducing viability of protoscoleces in CE patients when compared to ABZ treatment alone [85]. Combination therapies targeting both stages, metacestodes and protoscoleces, could also be assessed in our *in vitro* model, and could possibly reduce this risk and therefore receive more attention in future.

In addition to its application in the search for novel treatments against echinococcosis, the drug screening cascade presented here could be also applied in other cestode models such as *Taenia spp.*. The motility assay could be additionally extrapolated to several trematode species causing severe diseases wordwide, such as *Fasciola hepatica*, *Clonorchis sinensis*, *Ophisthorchis spp.*, or *Paragonimus spp.*, causing 90,000, 523,000, 188,000 and 1,049,000 DALYs, respectively [4]. Finally, it is important to keep in mind that more extensive and elaborate *in vitro* assessments of compound activities has a considerable potential to reduce the numbers of animal experiments, and is therefore highly relevant in terms of the 3R concept (reduce, replace, refine) [86].

## Acknowledgements

The authors thank Diana Gliga and Ursula Kurath for the DNA isolation and diagnostic confirmation of four *E. granulosus s.s.* isolates. We acknowledge the funding by the Swiss National Foundation that covered parts of this study (192072).

## Supporting information

**S1 Fig. Phylogenetic tree of *E. granulosus s.s.* isolates.** Four isolates of *E. granulosus s.s.* were genotyped using a concatenated sequence of the four mitochondrial markers *atp 6*, *nad I*, *cox I* and *rrnL*. The tree was generated according to the maximum likelihood method with the HKY+G+I model.

## References

[1] Eckert J, Deplazes P. Biological, epidemiological, and clinical aspects of echinococcosis, a zoonosis of increasing concern. Clin Microbiol Rev 2004;17:107–35.

[2] Budke CM, Deplazes P, Torgerson PR. Global Socioeconomic Impact of Cystic Echinococcosis. Emerg Infect Dis 2006;12:296–303. https://doi.org/10.3201/eid1202.050499.

[3] Bouwknegt M, Devleesschauwer B, Graham H, Robertson LJ, Giessen JW van der, null The Euro-Fbp workshop participants. Prioritisation of food-borne parasites in Europe, 2016. Euro Surveill Bull Eur Sur Mal Transm Eur Commun Dis Bull 2018;23. https://doi.org/10.2807/1560-7917.ES.2018.23.9.17-00161.

[4] Torgerson PR, Devleesschauwer B, Praet N, Speybroeck N, Willingham AL, Kasuga F, et al. World Health Organization Estimates of the Global and Regional Disease Burden of 11 Foodborne Parasitic Diseases, 2010: A Data Synthesis. PLoS Med 2015;12:e1001920. https://doi.org/10.1371/journal.pmed.1001920.

[5] FAO/WHO. Multicriteria-based ranking for risk management of food-borne parasites. 2014.

[6] Rossi P, Tamarozzi F, Galati F, Akhan O, Cretu CM, Vutova K, et al. The European Register of Cystic Echinococcosis, ERCE: state-of-the-art five years after its launch. Parasit Vectors 2020;13:236. https://doi.org/10.1186/s13071-020-04101-6.

[7] Rossi P, Tamarozzi F, Galati F, Pozio E, Akhan O, Cretu CM, et al. The first meeting of the European Register of Cystic Echinococcosis (ERCE). Parasit Vectors 2016;9:243. https://doi.org/10.1186/s13071-016-1532-3.

[8] Woolsey ID, Miller AL. Echinococcus granulosus sensu lato and Echinococcus multilocularis: A review. Res Vet Sci 2021;135:517–22. https://doi.org/10.1016/j.rvsc.2020.11.010.

[9] Gottstein B, Stojkovic M, Vuitton DA, Millon L, Marcinkute A, Deplazes P. Threat of alveolar echinococcosis to public health--a challenge for Europe. Trends Parasitol 2015;31:407–12. https://doi.org/10.1016/j.pt.2015.06.001.

[10] Trotz-Williams LA, Mercer NJ, Walters JM, Wallace D, Gottstein B, Osterman-Lind E, et al. Public Health Follow-up of Suspected Exposure to Echinococcus multilocularis in Southwestern Ontario. Zoonoses Public Health 2017;64:460–7. https://doi.org/10.1111/zph.12326.

[11] Galeh TM, Spotin A, Mahami-Oskouei M, Carmena D, Rahimi MT, Barac A, et al. The seroprevalence rate and population genetic structure of human cystic echinococcosis in the Middle East: A systematic review and meta-analysis. Int J Surg 2018;51:39–48. https://doi.org/10.1016/j.ijsu.2018.01.025.

[12] Hotez PJ. The rise or fall of neglected tropical diseases in East Asia Pacific. Acta Trop 2020;202:105182. https://doi.org/10.1016/j.actatropica.2019.105182.

[13] Torgerson PR. The emergence of echinococcosis in central Asia. Parasitology 2013;140:1667–73. https://doi.org/10.1017/S0031182013000516.

[14] Brehm K, Koziol U. Echinococcus–Host Interactions at Cellular and Molecular Levels. Adv. Parasitol., vol. 95, Elsevier; 2017, p. 147–212. https://doi.org/10.1016/bs.apar.2016.09.001.

[15] Díaz A, Casaravilla C, Irigoín F, Lin G, Previato JO, Ferreira F. Understanding the laminated layer of larval Echinococcus I: structure. Trends Parasitol 2011;27:204–13. https://doi.org/10.1016/j.pt.2010.12.012.

[16] Koziol U, Rauschendorfer T, Zanon Rodríguez L, Krohne G, Brehm K. The unique stem cell system of the immortal larva of the human parasite Echinococcus multilocularis. EvoDevo 2014;5:10. https://doi.org/10.1186/2041-9139-5-10.

[17] Koziol U, Krohne G, Brehm K. Anatomy and development of the larval nervous system in Echinococcus multilocularis. Front Zool 2013;10:24. https://doi.org/10.1186/1742-9994-10-24.

[18] Brunetti E, Kern P, Vuitton DA. Expert consensus for the diagnosis and treatment of cystic and alveolar echinococcosis in humans. Acta Trop 2010;114:1–16. https://doi.org/10.1016/j.actatropica.2009.11.001.

[19] Grüner B, Kern P, Mayer B, Gräter T, Hillenbrand A, Barth TFE, et al. Comprehensive diagnosis and treatment of alveolar echinococcosis: A single-center, long-term observational study of 312 patients in Germany. GMS Infect Dis 2017:1–12. https://doi.org/doi:10.3205/id000027.

[20] Reuter S, Buck A, Manfras B, Kratzer W, Seitz HM, Darge K, et al. Structured treatment interruption in patients with alveolar echinococcosis. Hepatol Baltim Md 2004;39:509–17. https://doi.org/10.1002/hep.20078.

[21] Schubert A, Koziol U, Cailliau K, Vanderstraete M, Dissous C, Brehm K. Targeting Echinococcus multilocularis stem cells by inhibition of the Polo-like kinase EmPlk1. PLoS Negl Trop Dis 2014;8:e2870. https://doi.org/10.1371/journal.pntd.0002870.

[22] Brehm K, Koziol U. On the importance of targeting parasite stem cells in anti-echinococcosis drug development. Parasite Paris Fr 2014;21:72. https://doi.org/10.1051/parasite/2014070.

[23] Ammann RW, Eckert J. Cestodes. Echinococcus. Gastroenterol Clin North Am 1996;25:655–89.

[24] Lundström-Stadelmann B, Rufener R, Ritler D, Zurbriggen R, Hemphill A. The importance of being parasiticidal… an update on drug development for the treatment of alveolar echinococcosis. Food Waterborne Parasitol 2019;15:e00040. https://doi.org/10.1016/j.fawpar.2019.e00040.

[25] Spiliotis M, Lechner S, Tappe D, Scheller C, Krohne G, Brehm K. Transient transfection of Echinococcus multilocularis primary cells and complete in vitro regeneration of metacestode vesicles. Int J Parasitol 2008;38:1025–39. https://doi.org/10.1016/j.ijpara.2007.11.002.

[26] Spiliotis M, Mizukami C, Oku Y, Kiss F, Brehm K, Gottstein B. Echinococcus multilocularis primary cells: improved isolation, small-scale cultivation and RNA interference. Mol Biochem Parasitol 2010;174:83–7. https://doi.org/10.1016/j.molbiopara.2010.07.001.

[27] Spiliotis M, Tappe D, Sesterhenn L, Brehm K. Long-term in vitro cultivation of Echinococcus multilocularis metacestodes under axenic conditions. Parasitol Res 2004;92:430–2. https://doi.org/10.1007/s00436-003-1046-8.

[28] Spiliotis M, Brehm K. Axenic in vitro cultivation of Echinococcus multilocularis metacestode vesicles and the generation of primary cell cultures. Methods Mol Biol Clifton NJ 2009;470:245–62. https://doi.org/10.1007/978-1-59745-204-5_17.

[29] Laurimäe T, Kronenberg PA, Rojas CAA, Ramp TW, Eckert J, Deplazes P. Long-term (35 years) cryopreservation of Echinococcus multilocularis metacestodes. Parasitology 2020;147:1048–54. https://doi.org/10.1017/S003118202000075X.

[30] Hemphill A, Lundström-Stadelmann B. Echinococcus: the model cestode parasite. Parasitology 2021;148:1401–5. https://doi.org/10.1017/S003118202100113X.

[31] Ritler D, Rufener R, Sager H, Bouvier J, Hemphill A, Lundström-Stadelmann B. Development of a movement-based in vitro screening assay for the identification of new anti-cestodal compounds. PLoS Negl Trop Dis 2017;11. https://doi.org/10.1371/journal.pntd.0005618.

[32] Stadelmann B, Rufener R, Aeschbacher D, Spiliotis M, Gottstein B, Hemphill A. Screening of the Open Source Malaria Box Reveals an Early Lead Compound for the Treatment of Alveolar Echinococcosis. PLoS Negl Trop Dis 2016;10. https://doi.org/10.1371/journal.pntd.0004535.

[33] Stadelmann B, Scholl S, Müller J, Hemphill A. Application of an in vitro drug screening assay based on the release of phosphoglucose isomerase to determine the structure-activity relationship of thiazolides against Echinococcus multilocularis metacestodes. J Antimicrob Chemother 2010;65:512–9. https://doi.org/10.1093/jac/dkp490.

[34] Eckert J, Weltgesundheitsorganisation, International Office of Epizootics, editors. WHO/OIE manual on Echinococcosis in humans and animals: a public health problem of global concern. Paris: World Organisation for Animal Health; 2001.

[35] WHO-Informal Working Group on Echinococcosis. Guidelines for treatment of cystic and alveolar echinococcosis in humans. Bull World Health Organ 1996;74:231–42.

[36] Horton RJ. Albendazole in treatment of human cystic echinococcosis: 12 years of experience. Acta Trop 1997;64:79–93. https://doi.org/10.1016/S0001-706X(96)00640-7.

[37] Bouaziz S, Amri M, Taibi N, Zeghir-Bouteldja R, Benkhaled A, Mezioug D, et al. Protoscolicidal activity of Atriplex halimus leaves extract against Echinococcus granulosus protoscoleces. Exp Parasitol 2021;229:108155. https://doi.org/10.1016/j.exppara.2021.108155.

[38] Mahmoudvand H, Khalaf AK, Beyranvand M. In Vitro and Ex Vivo Evaluation of Capparis spinosa Extract to Inactivate Protoscoleces During Hydatid Cyst Surgery. Curr Drug Discov Technol 2021;18:e18082020185049. https://doi.org/10.2174/1570163817999200819091336.

[39] Muhedier M, Li J, Liu H, Ma G, Amahong K, Lin R, et al. Tacrolimus, a rapamycin target protein inhibitor, exerts anti-cystic echinococcosis effects both in vitro and in vivo. Acta Trop 2020;212:105708. https://doi.org/10.1016/j.actatropica.2020.105708.

[40] Wen L, Lv G, Zhao J, Lu S, Gong Y, Li Y, et al. In vitro and in vivo Effects of Artesunate on Echinococcus granulosus Protoscoleces and Metacestodes. Drug Des Devel Ther 2020;14:4685–94. https://doi.org/10.2147/DDDT.S254166.

[41] Casado N, Pérez-Serrano J, Denegri G, Rodríguez-Caabeiro F. Development of a chemotherapeutic model for the in vitro screening of drugs against Echinococcus granulosus cysts: the effects of an albendazole-albendazole sulphoxide combination. Int J Parasitol 1996;26:59–65. https://doi.org/10.1016/0020-7519(95)00095-X.

[42] Elissondo M, Ceballos L, Dopchiz M, Andresiuk V, Alvarez L, Bruni SS, et al. In vitro and in vivo effects of flubendazole on Echinococcus granulosus metacestodes. Parasitol Res 2007;100:1003–9. https://doi.org/10.1007/s00436-006-0381-y.

[43] Walker M, Rossignol JF, Torgerson P, Hemphill A. In vitro effects of nitazoxanide on Echinococcus granulosus protoscoleces and metacestodes. J Antimicrob Chemother 2004;54:609–16. https://doi.org/10.1093/jac/dkh386.

[44] Albani CM, Elissondo MC, Cumino AC, Chisari A, Denegri GM. Primary cell culture of Echinococcus granulosus developed from the cystic germinal layer: biological and functional characterization. Int J Parasitol 2010;40:1269–75. https://doi.org/10.1016/j.ijpara.2010.03.008.

[45] Albani CM, Cumino AC, Elissondo MC, Denegri GM. Development of a cell line from Echinococcus granulosus germinal layer. Acta Trop 2013;128:124–9. https://doi.org/10.1016/j.actatropica.2013.07.001.

[46] Fiori PL, Monaco G, Scappaticci S, Pugliese A, Canu N, Cappuccinelli P. Establishment of cell cultures from hydatid cysts of Echinococcus granulosus. Int J Parasitol 1988;18:297–305. https://doi.org/10.1016/0020-7519(88)90137-3.

[47] Echeverría CI, Isolabella DM, Gonzalez EAP, Leonardelli A, Prada L, Perrone A, et al. Morphological and biological characterization of cell line developed from bovine Echinococcus granulosus. Vitro Cell Dev Biol -Anim 2010;46:781–92. https://doi.org/10.1007/s11626-010-9345-8.

[48] Thompson RCA. Biology and Systematics of Echinococcus. Adv Parasitol 2017;95:65–109. https://doi.org/10.1016/bs.apar.2016.07.001.

[49] Pensel PE, Albani C, Gamboa GU, Benoit JP, Elissondo MC. In vitro effect of 5-fluorouracil and paclitaxel on Echinococcus granulosus larvae and cells. Acta Trop 2014;140:1–9. https://doi.org/10.1016/j.actatropica.2014.07.013.

[50] Pensel PE, Elissondo N, Gambino G, Gamboa GU, Benoit JP, Elissondo MC. Experimental cystic echinococcosis therapy: In vitro and in vivo combined 5-fluorouracil/albendazole treatment. Vet Parasitol 2017;245:62–70. https://doi.org/10.1016/j.vetpar.2017.08.011.

[51] Marinova I, Spiliotis M, Wang J, Muhtarov M, Chaligiannis I, Sotiraki S, et al. Molecular characterization of Echinococcus granulosus isolates from Bulgarian human cystic echinococcosis patients. Parasitol Res 2017;116:1043–54. https://doi.org/10.1007/s00436-017-5386-1.

[52] Nakao M, Sako Y, Ito A. The Mitochondrial Genome of the Tapeworm Taenia solium: A Finding of the Abbreviated Stop Codon U. J Parasitol 2003;89:633–5. https://doi.org/10.1645/0022-3395(2003)089[0633:TMGOTT]2.0.CO;2.

[53] Nakao M, Yokoyama N, Sako Y, Fukunaga M, Ito A. The complete mitochondrial DNA sequence of the cestode Echinococcus multilocularis (Cyclophyllidea: Taeniidae). Mitochondrion 2002;1:497–509. https://doi.org/10.1016/S1567-7249(02)00040-5.

[54] Nakao M, McMANUS DP, Schantz PM, Craig PS, Ito A. A molecular phylogeny of the genus Echinococcus inferred from complete mitochondrial genomes. Parasitology 2006;134:713–22. https://doi.org/10.1017/S0031182006001934.

[55] Le TH, Pearson MS, Blair D, Dai N, Zhang LH, Mcmanus DP. Complete mitochondrial genomes confirm the distinctiveness of the horse-dog and sheep-dog strains of Echinococcus granulosus. Parasitology 2002;124:97–112. https://doi.org/10.1017/S0031182001008976.

[56] Wang N, Xie Y, Liu T, Zhong X, Wang J, Hu D, et al. The complete mitochondrial genome of G3 genotype of Echinococcus granulosus (Cestoda: Taeniidae). Mitochondrial DNA Part A 2016;27:1701–2. https://doi.org/10.3109/19401736.2014.961129.

[57] Nakao M, Yanagida T, Konyaev S, Lavikainen A, Odnokurtsev VA, Zaikov VA, et al. Mitochondrial phylogeny of the genus Echinococcus (Cestoda: Taeniidae) with emphasis on relationships among Echinococcus canadensis genotypes. Parasitology 2013;140:1625–36. https://doi.org/10.1017/S0031182013000565.

[58] Tamura K, Stecher G, Kumar S. MEGA11: Molecular Evolutionary Genetics Analysis Version 11. Mol Biol Evol 2021;38:3022–7. https://doi.org/10.1093/molbev/msab120.

[59] Mäser P, Grether-Bühler Y, Kaminsky R, Brun R. An anti-contamination cocktail for the in vitro isolation and cultivation of parasitic protozoa. Parasitol Res 2002;88:172–4. https://doi.org/10.1007/s00436-001-0511-5.

[60] Herz M, Brehm K. Serotonin stimulates Echinococcus multilocularis larval development. Parasit Vectors 2021;14:14. https://doi.org/10.1186/s13071-020-04533-0.

[61] Rufener R, Dick L, D’Ascoli L, Ritler D, Hizem A, Wells TNC, et al. Repurposing of an old drug: *In vitro* and *in vivo* efficacies of buparvaquone against *Echinococcus multilocularis*. Int J Parasitol Drugs Drug Resist 2018;8:440–50. https://doi.org/10.1016/j.ijpddr.2018.10.011.

[62] Lundström-Stadelmann B, Rufener R, Hemphill A. Drug repurposing applied: Activity of the anti-malarial mefloquine against Echinococcus multilocularis. Int J Parasitol Drugs Drug Resist 2020;13:121–9. https://doi.org/10.1016/j.ijpddr.2020.06.002.

[63] Rufener R, Ritler D, Zielinski J, Dick L, da Silva ET, da Silva Araujo A, et al. Activity of mefloquine and mefloquine derivatives against Echinococcus multilocularis. Int J Parasitol Drugs Drug Resist 2018;8:331–40. https://doi.org/10.1016/j.ijpddr.2018.06.004.

[64] Bryant C. Electron transport in parasitic helminths and protozoa. Adv Parasitol 1970;8:139–72.

[65] Spicher M, Roethlisberger C, Lany C, Stadelmann B, Keiser J, Ortega-Mora LM, et al. In Vitro and In Vivo Treatments of Echinococcus Protoscoleces and Metacestodes with Artemisinin and Artemisinin Derivatives. Antimicrob Agents Chemother 2008;52:3447–50. https://doi.org/10.1128/AAC.00553-08.

[66] Karpstein T, Chaudhry S, Bresson-Hadni S, Hayoz M, Boubaker G, Hemphill A, et al. Maca against Echinococcosis?-A Reverse Approach from Patient to In Vitro Testing. Pathog Basel Switz 2021;10:1335. https://doi.org/10.3390/pathogens10101335.

[67] Mehmood N, Dessì G, Ahmed F, Joanny G, Tamponi C, Cappai MG, et al. Genetic diversity and transmission patterns of Echinococcus granulosus sensu stricto among domestic ungulates of Sardinia, Italy. Parasitol Res 2021;120:2533–42. https://doi.org/10.1007/s00436-021-07186-9.

[68] Varcasia A, Canu S, Lightowlers MW, Scala A, Garippa G. Molecular characterization of Echinococcus granulosus strains in Sardinia. Parasitol Res 2006;98:273–7. https://doi.org/10.1007/s00436-005-0059-x.

[69] Alvarez Rojas CA, Romig T, Lightowlers MW. Echinococcus granulosus sensu lato genotypes infecting humans--review of current knowledge. Int J Parasitol 2014;44:9–18. https://doi.org/10.1016/j.ijpara.2013.08.008.

[70] Rodriguez-Caabeiro F, Casado N. Evidence of in vitro germinal layer development in Echinococcus granulosus cysts. Parasitol Res 1988;74:558–62. https://doi.org/10.1007/BF00531634.

[71] Dezaki ES, Yaghoubi MM, Spiliotis M, Boubaker G, Taheri E, Almani PG, et al. Comparison of ex vivo harvested and in vitro cultured materials from Echinococcus granulosus by measuring expression levels of five genes putatively involved in the development and maturation of adult worms. Parasitol Res 2016;115:4405–16. https://doi.org/10.1007/s00436-016-5228-6.

[72] Chaudhry S, Zurbriggen R, Preza M, Kämpfer T, Kaethner M, Memedovski R, et al. Dual inhibition of the Echinococcus multilocularis energy metabolism. Front Vet Sci 2022;9:981664. https://doi.org/10.3389/fvets.2022.981664.

[73] Hemphill A, Stadelmann B, Rufener R, Spiliotis M, Boubaker G, Müller J, et al. Treatment of echinococcosis: albendazole and mebendazole – what else? Parasite 2014;21. https://doi.org/10.1051/parasite/2014073.

[74] Ingold K, Bigler P, Thormann W, Cavaliero T, Gottstein B, Hemphill A. Efficacies of albendazole sulfoxide and albendazole sulfone against In vitro-cultivated Echinococcus multilocularis metacestodes. Antimicrob Agents Chemother 1999;43:1052–61.

[75] Küster T, Stadelmann B, Aeschbacher D, Hemphill A. Activities of fenbendazole in comparison with albendazole against Echinococcus multilocularis metacestodes in vitro and in a murine infection model. Int J Antimicrob Agents 2014;43:335–42. https://doi.org/10.1016/j.ijantimicag.2014.01.013.

[76] Ali-Khan Z, Siboo R. Pathogenesis and host response in subcutaneous alveolar hydatidosis. I. Histogenesis of alveolar cyst and a qualitative analysis of the inflammatory infiltrates. Z Für Parasitenkd Berl Ger 1980;62:241–54.

[77] Hinz E. Die Aufbereitung des Infektionsmaterials für die intraperitoneale Infektion der Maus mit Echinococcus multilocularis. Z FÜR TROPENMEDIZIN Parasitol 1973;4:387–90.

[78] Küster T, Hermann C, Hemphill A, Gottstein B, Spiliotis M. Subcutaneous infection model facilitates treatment assessment of secondary Alveolar echinococcosis in mice. PLoS Negl Trop Dis 2013;7:e2235. https://doi.org/10.1371/journal.pntd.0002235.

[79] Liance M, Vuitton DA, Guerret-Stocker S, Carbillet JP, Grimaud JA, Houin R. Experimental alveolar echinococcosis. Suitability of a murine model of intrahepatic infection by Echinococcus multilocularis for immunological studies. Experientia 1984;40:1436–9.

[80] Ohbayashi M. STUDIES ON ECHINOCOCCOSIS X. : HISTOLOGICAL OBSERVATIONS ON EXPERIMENTAL CASES OF MULTILOCULAR ECHINOCOCCOSIS. Jpn J Vet Res 1960;8:134–60.

[81] Yamashita J, Ohbayashi M, Konno S. Studies on echinococcosis VI. Secondary Echinococcus multilocularis in mice. Jpn J Vet Res 1957:197–202.

[82] Dempster RP, Berridge MV, Harrison GBL, Heath DD. Echinococcus granulosus: Development of an intermediate host mouse model for use in vaccination studies. Int J Parasitol 1991;21:549–54. https://doi.org/10.1016/0020-7519(91)90059-G.

[83] Li Z, Zhang C, Li L, Bi X, Li L, Yang S, et al. The local immune response during Echinococcus granulosus growth in a quantitative hepatic experimental model. Sci Rep 2019;9:19612. https://doi.org/10.1038/s41598-019-56098-3.

[84] Zhang W, You H, Zhang Z, Turson G, Hasyet A, McManus DP. Further studies on an intermediate host murine model showing that a primary Echinococcus granulosus infection is protective against subsequent oncospheral challenge. Parasitol Int 2001;50:279–83. https://doi.org/10.1016/S1383-5769(01)00086-1.

[85] Cobo F, Yarnoz C, Sesma B, Fraile P, Aizcorbe M, Trujillo R, et al. Albendazole plus praziquantel versus albendazole alone as a pre-operative treatment in intra-abdominal hydatisosis caused by Echinococcus granulosus. Trop Med Int Health TM IH 1998;3:462–6.

[86] Russell WMS, Burch RL. The principles of Humane Experimental Technique. Methuen Lond 1959.

